# BRWD1 orchestrates chromatin topology by converting static to dynamic cohesin complexes

**DOI:** 10.1101/2023.01.23.525212

**Authors:** Malay Mandal, Mark Maienschein-Cline, Yeguang Hu, Azam Mohsin, Margaret L. Veselits, Nathaniel E. Wright, Michael K. Okoreeh, Young me Yoon, Jacob Veselits, Katia Georgopoulos, Marcus R. Clark

## Abstract

Lymphocyte development consists of sequential and mutually exclusive cell states of proliferative selection and antigen receptor gene recombination^1^. Transitions between each state require large, coordinated changes in epigenetic landscapes and transcriptional programs^2,3^. How this occurs remains unclear. Herein, we demonstrate that in small pre-B cells, the lineage and stagespecific epigenetic reader Bromodomain and WD Repeating Containing Protein 1 (BRWD1)^2,4^ reorders three-dimensional chromatin topology to affect transition between proliferative and gene recombination molecular programs. BRWD1 regulated the switch between poised and active enhancers interacting with promoters and coordinated this with *Igk* locus contraction. BRWD1 did so by converting chromatin-bound static cohesin to dynamic complexes competent to mediate long-range looping. Remarkably, ATP depletion recapitulated cohesin distributions observed in *Brwd1^-/-^* cells. Therefore, in small pre-B cells, cohesin conversion is the main energetic mechanism dictating where dynamic looping occurs in the genome. Our findings provide a new mechanism of cohesin regulation and reveal how cohesin function can be dictated by lineage contextual mechanisms to facilitate specific cell fate transitions.

## Main

### BRWD1 orchestrates chromatin topology

Developmental transit between proliferating large pre-B cells and small pre-B cells initiating *Igk* recombination requires a profound switch in genetic program including differential expression of over 7,000 genes and changes in accessibility at over 50,000 genomic sites^2^. Transit from the proliferative to recombinatorial programs is mediated by BRWD1, which represses early developmental enhancers and opens those of late B cell development^2,4^. BRWD1 is also required for recombination at *Igk* where it opens Jk for assembly of the recombination complex^4^. BRWD1 is repressed by IL-7 receptor signaling and induced by pre-BCR and CXCR4 signaling which also dictate the epigenetic landscape [histone 3 (H3), lysine 9 acetylation (K9Ac), serine S10 phosphorylation (S10p) and K14Ac] to which BRWD1 is recruited^4,5^. This coordinate regulation ensures that BRWD1 first functions in small pre-B cells. Interestingly, BRWD1 regulates accessibility of enhancers distant from BRWD1 chromatin binding sites^2^. We hypothesized that BRWD1 does so either through existing three-dimensional (3D) chromatin interactions or by regulating such interactions.

Therefore, we performed *in situ* Hi-C on wild-type (WT) and *Brwd1^-/-^* small pre-B cells, and WT immature B cells^6^. We used WT, as opposed to small pre-B cells expressing a mutant *Rag1* and transgenic immunoglobulin heavy chain (*Rag1^-/-^B1-8i*), as the latter are aberrant intermediates between WT large pre-B and small pre-B cells (Extended Data Fig.1a)^7^. Furthermore, our gating strategy for WT small pre-B cells captures cells in which *Igk* is actively transcribed yet minimially recombined^8^. Principal component analysis (PCA) demonstrated that each replicate was similar, and that each population had different 3D chromatin topologies (Fig. 1a). All populations had features of committed B cell progenitors with well-formed topological associating domains (TADs) at the B cell specific *Ebf1* gene but not at the *Bcl11b* (T cells) or *Il1f9* (granulocytes) genes (Extended Data Fig. 1b)^9–11^.

**Figure 1.**
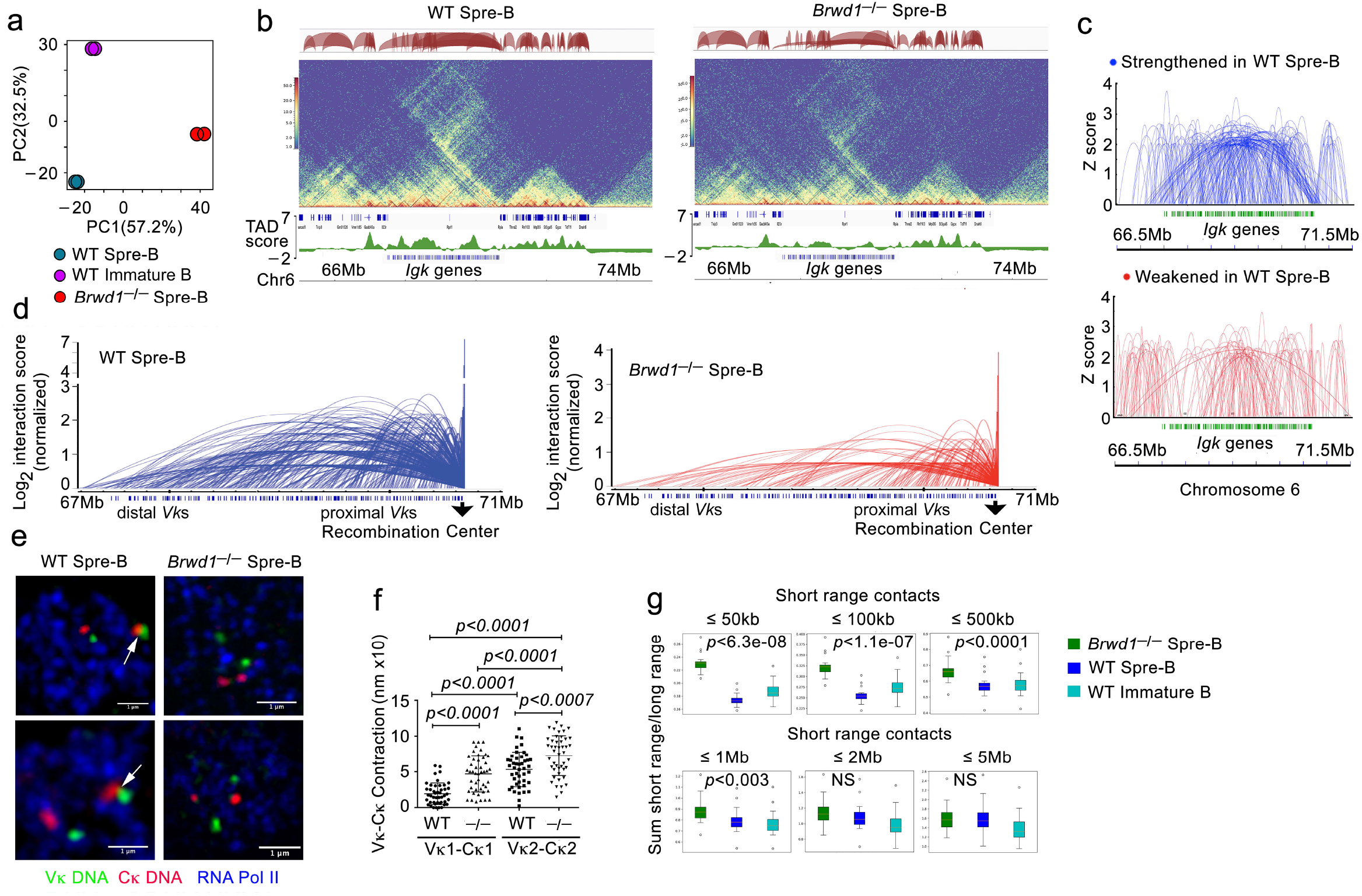
BRWD1 enhances long-range genomic interactions including *Igk* locus contraction. **a,** Principal Component Analysis (PCA) of Hi-C data obtained from indicated flow-purified cell populations. **b,** Chromatin interactions at the *Igk* locus (top) in WT (left) and *Brwd1^-/-^* (right) small pre-B cells. Height of the loops indicate the strength of the loop. Corresponding Hi-C contact maps (10 kb resolution, middle) and TAD scores (bottom) are shown. **c,** Arc plots indicating strengthened and weakened interactions (*P*<0.05) across the *Igk* locus in WT compared to *Brwd1^-/-^* small pre-B cells. **d,** Point-of-view arc plots demonstrating normalized genomic interactions between the recombination center (*Jk-Ck*) and all *Vk* gene segments in WT (left) and *Brwd1^-/-^* (right) small pre-B cells. **e,** *Vk* and *Ck* DNA-FISH in combination with elongating RNA Pol II immuno-confocal microscopy on WT and *Brwd1^-/-^* small pre-B cells. Shown are representative images (40 cells/sample; n = 2 experiments). Each example demonstrates a subnuclear field of view. Scale bar, 1μm. **f**, Quantification of genomic contraction for both *Igk* alleles in WT and *Brwd1^-/-^* small pre-B cells. **g,** Box plots showing sum of short over long-range genomic interactions in indicated cell populations. Lengths considered to be short-range interactions, versus all longer range interactions, are indicated. Resolution: 5kb. *P* values were determined by Wilcoxon test.

BRWD1 is required for *Igk* recombination^4^. Plotting interactions at *Igk* (~3 mb, Fig 1b) that were either strengthened or weakened in WT small pre-B cells revealed that BRWD1 enhanced long-range interactions and repressed short range interactions (Fig. 1c). Looking from the point of view of the *Jk* gene region, where the recombination center assembles^4,12^, BRWD1 enhanced interactions with both proximal and distal *Vk* regions (Fig. 1d). Consistent with these observations, immuno-FISH using a DNA probe for distal *Vk* gene segments, and one for the *Ck* region, confirmed that BRWD1 contributed to, or maintained, locus contraction in small pre-B cells (Fig. 1e,f)^13^.

BRWD1 represses *Myc* transcription, and this is associated with silencing of distal enhancers^2^. In *Brwd1^-/-^* small pre-B cells, there was enhanced looping from the *Myc* promoter to regions across the *Myc* locus (~0.8 Mb) (Extended data Fig. 2a) including specific distal enhancers (Extended Data Fig. 2b). BRWD1 also repressed looping and expression at the *Mki67* and *H2-Q1* loci (Extended Data Fig. 2c,d). However, genome-wide analyses revealed that the overall effect of BRWD1 was to induce long-range looping, and repress short-range looping, over distances of up to 1 Mb (Fig. 1g).

### BRWD1 regulates enhancer transitions

As BRWD1 regulates enhancer accessibility^2^, we further examined enhancer state by performing chromatin immunoprecipitation followed by sequencing (ChIP-seq) for the epigenetic marks H3K4me1, which identifies all enhancers, and H3K27Ac which identifies those that are activated^14^. Both marks were reproducibly detected across the genome (Extended Data Fig. 3a). However, the prevalence and distributions of these marks were significantly different between WT and *Brwd1^-/-^* small pre-B cells (Fig. 2a). The overall number of H3K4me1 peaks was similar, while approximately 50k H3K27Ac peaks were lost in *Brwd1^-/-^* small pre-B cells. However, only 61% of H3K4me1 peaks and approximately 63% of H3K27Ac peaks were shared between WT and *Brwd1^-/-^* small pre-B cells. In addition to H3K4me1 and H3K27Ac, active enhancers are more accessible (assay for transposase-accessible chromatin sequencing, ATAC-seq) than poised or repressed enhancers^15,16^. When active enhancers were defined as open regions containg both H3K4me1 and H3K27Ac, there were 17,244 active enhancer peaks in WT and 35,481 in *Brwd1^-/-^* small pre-B cells (Fig. 2b). The distribution of active enhancers in the two cell populations were different with 10,862 enhancer peaks specific to WT and 18,561 enhancer peaks specific to *Brwd1^-/-^* small pre-B cells (Fig. 2c). Of active enhancer sites specific to WT small pre-B cells, the main differential epigenetic mark was H3K27Ac, while H3K4me1 peak intensity distributions were similar (Fig. 2d). The same pattern was observed at those active enhancer peaks specific to *Brwd1^-/-^* cells.

**Figure 2.**
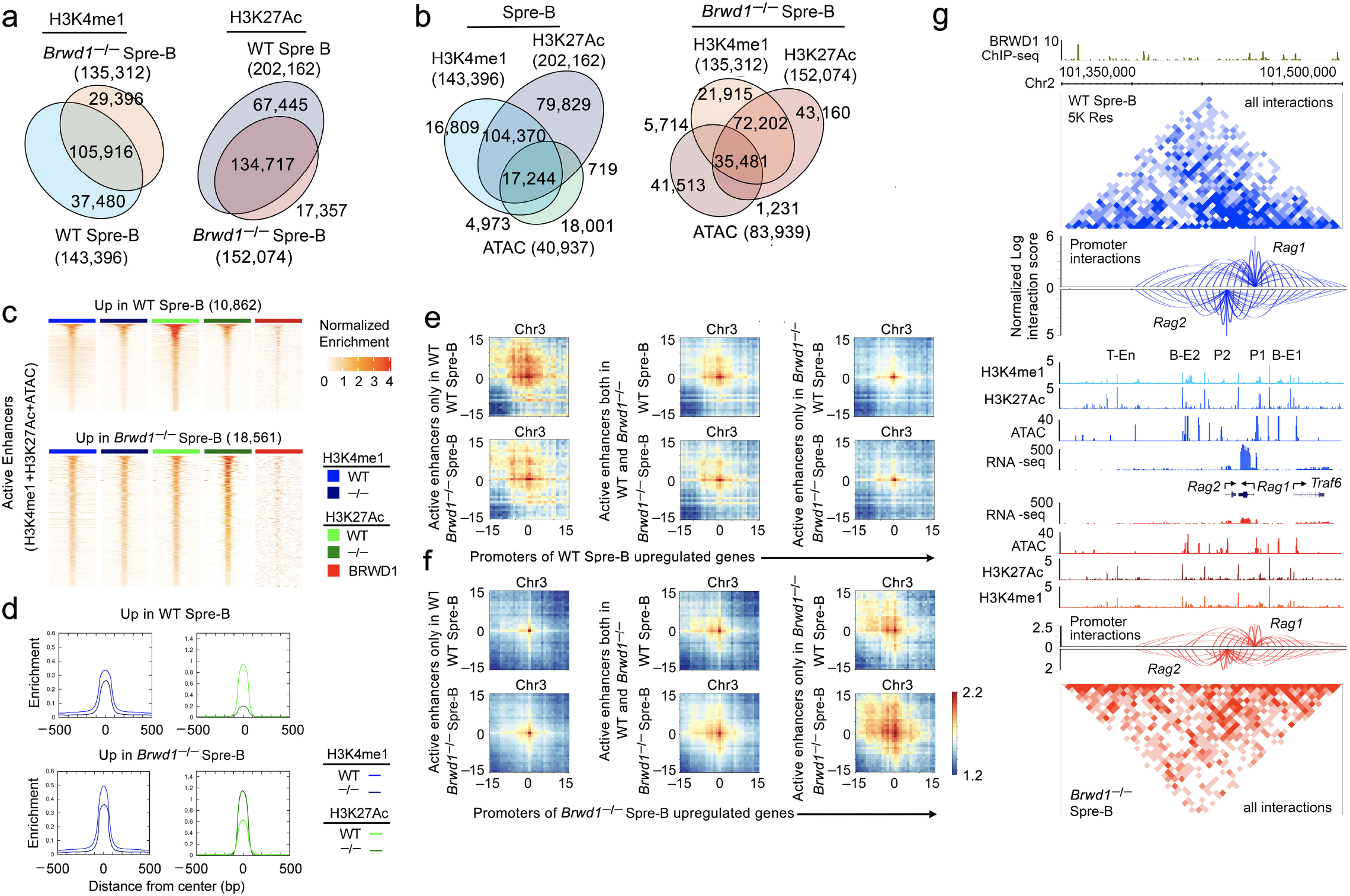
BRWD1 regulates transition from poised to active enhancers interacting with promoters. **a,** Total and coincident H3K4me1 and H3K27Ac peaks determined by ChIP-seq in WT and *Brwd1^-/-^* small pre-B cells. Total number of peaks for each population is shown in parentheses with the number in each Venn region indicated. **b,** Total and coincident H3K4me1 and H3K27Ac peaks determined by ChIP-seq and overlapping open chromatin peaks (ATAC-seq) in indicated B-cell progenitor populations. **c,** Heatmap of tag density of H3K4me1, H3K27Ac and BRWD1 chromatin binding at differentially regulated enhancers in WT and *Brwd1^-/-^* small pre-B cells. Binding is plotted ±500 base pairs from active enhancer centers (H3K4me1+H3K27Ac+ATAC) upregulated in WT (top) and *Brwd1^-/-^* (bottom) small pre-B cells. Differential peaks called by FDR<0.05 and logFC>1. **d,** Histograms of H3K4me1 and H3K27Ac enrichment at active enhancers upregulated in WT and *Brwd1^-/-^* small pre-B cells. **e-f,** Hi-C aggregate plots of contacts between promoters and active enhancers for chromosome 3. **e**, Promoters of genes upregulated in WT and **f**, those upregulated in *Brwd1^-/-^* small pre-B cells. Left panels show active enhancers only in WT small pre-B cells, middle panels show common active enhancers and right panels show active enhancers only in *Brwd1^-/-^* small pre-B cells. Differential observed/expected Hi-C signals were plotted. Contacts centered at all indicated promoters and enhancers on chromosome 3. Plots show both intensity (color) and breadth of Hi-C interactions between promoters and enhancers. **g,** *Rag1-Rag2* gene region demonstrating BRWD1 chromatin binding, Hi-C contact matrices at 5 kb resolution, point of view arc plots from the *Rag1* and *Rag2* promoters, indicated ChIP-seq, ATAC-seq and RNA-seq for WT (upper blue panels) and *Brwd1^-/-^* small pre-B cells (lower red panels). The T cell specific (T-En) and B cell specific (B-E1 and B-E2) enhancers are indicated as well as the *Rag1* (P1) and *Rag2* (P2) promoters.

Super-enhancers (SEs) can be defined as regions with high density H3K27Ac over extended distances^17^ (Extended Data Fig. 3b). The number of super-enhancers in WT and *Brwd1^-/-^* cells were similar (Extended Data Fig. 3c). However, only approximately 60% of super-enhancers in WT small pre-B cells and 62% of those in *Brwd1^-/-^* small pre-B cells were shared. As evident at other enhancers, BRWD1 primarily regulated the genomic prevalence of H3K27Ac, and not H3K4me1, peaks (Extended data Fig. 3d-g).

However, the preferential regulation of H3K27Ac by BRWD1 is not universal to all genomic sites. For example, at *Igk*, BRWD1 is necessary for also imparting H3K4me1 throughout the Jk to 3’k enhancer (3’Ek) region (Extended data Fig. 3h). This suggests that at *Igk*, BRWD1 orchestrates a more complex regulation than at most other genomic loci.

We then examined the relationship between enhancer activation and their interactions with promoters. As can be seen at a representative chromosome, the intensity and breadth of genomic interactions correlated with both enhancer activation state and transcription at promoters (Fig. 2e-f). Specific examples illustrate how BRWD1 coordinately regulates H3K27Ac marks, enhancer accessibility, chromatin 3D topology, and gene transcription (Fig. 2g and Extended Data Fig. 4a-c). These data indicate that BRWD1 controls small pre-B cell specific developmental transitions between poised enhancers marked with H3K4me1 and active enhancers that interact with promoters and induce transcription.

### BRWD1 regulates distribution of chromatin-bound cohesin complexes

Cohesin is a multimeric complex that mediates chromatin extrusion and has been implicated in promoter-enhancer interactions and immunoglobulin gene contraction^18–26^. Expression of most cohesin components was higher in *Brwd1^-/-^* small pre-B cells, except *Nipbl* (1.3x lower) (Extended Data Fig.5a), the protein product of which both loads the cohesin complex and enhances loop extrusion processivity^19,27–29^. We next performed ChIP-seq for CTCF, the cohesin ring components RAD21 and SMC3, NIPBL and WAPL. The latter mediates both cohesin chromatin release and recycling^30,31^. The number of peaks for CTCF and each cohesin component was similar between WT and *Brwd1^-/-^* small pre-B cells (Fig. 3a). Furthermore, the distribution of CTCF peaks was similar (Fig. 3b). In contrast, distributions of cohesin chromatin subunit binding were very different with only a minority of total subunit peaks conserved between WT and *Brwd1^-/-^* small pre-B cells (Fig. 3c). Furthermore, distributions of relative binding intensities were different in *Brwd1^-/-^* small pre-B cells (Fig. 3d-g and Extended Data Fig. 5b-i). While there was a single linear intensity relationship between SMC3 and RAD21 in WT small pre-B cells, there were two discrete binding complexes in *Brwd1^-/-^* cells (Fig. 3d). In contrast, in WT cells, there were two distinct cohesin complexes of NIPBL with either RAD21 or SMC3, but only one in *Brwd1^-/-^* cells (Fig. 3e,f). Furthemore, there was a shift in binding intensity of WAPL with RAD21 in *Brwd1^-/-^* cells (Fig. 3g). These findings suggest a general reordering of cohesin subunit chromatin binding in *Brwd1^-/-^* small pre-B cells.

**Figure 3.**
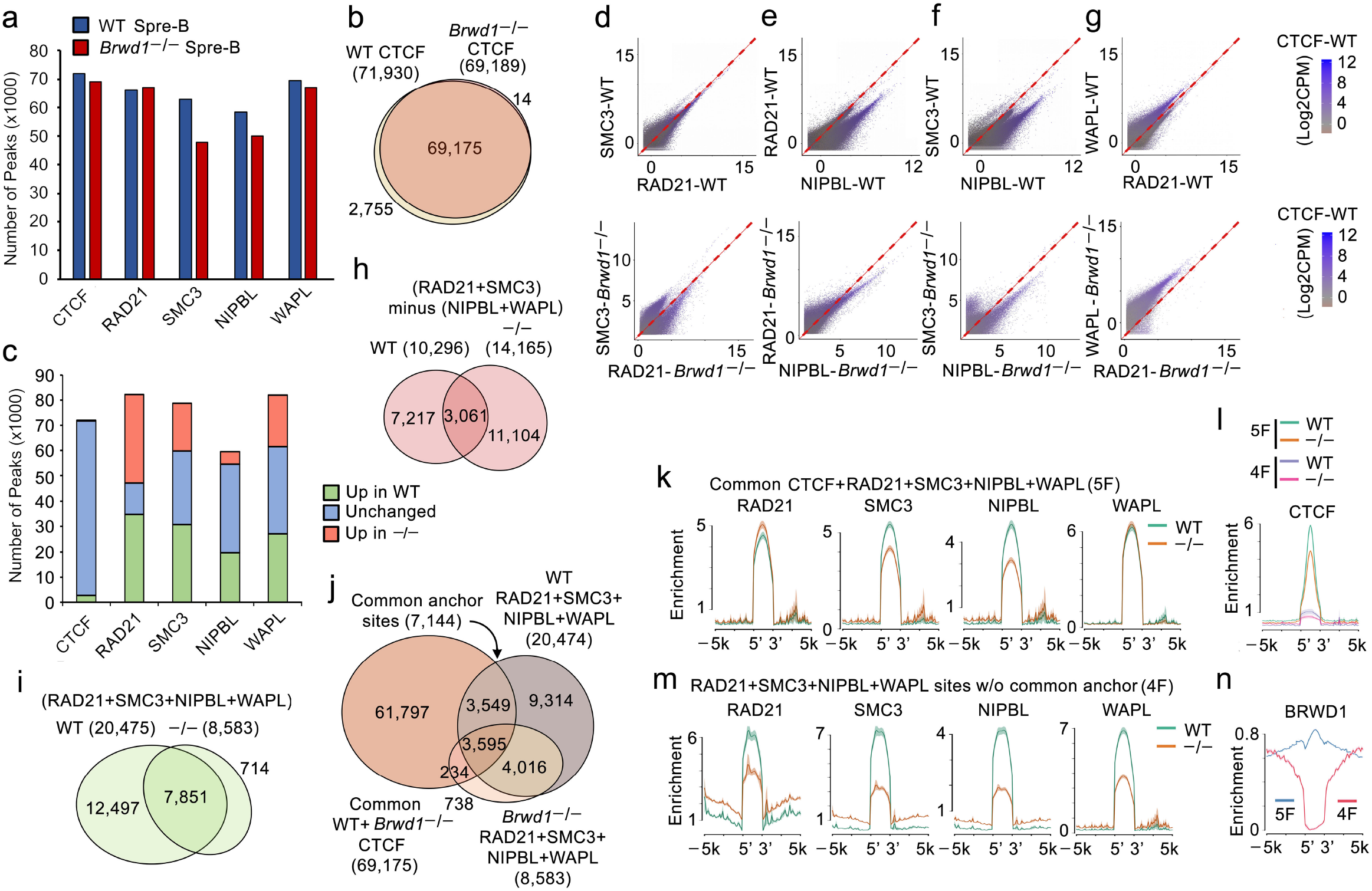
BRWD1 modulates chromatin binding of different cohesin complexes. **a**, Bar plot of the number of ChIP-seq peaks for CTCF, RAD21, SMC3, NIPBL and WAPL in WT and *Brwd1^-/-^* small pre-B cells. **b,** Total and coincident CTCF binding peaks in WT and *Brwd1^-/-^* small pre-B cells. **c,** Number of ChIP-seq peaks of CTCF, RAD21, SMC3, NIPBL and WAPL differentially enriched in WT and *Brwd1^-/-^* small pre-B cells. **d-g,** Scatterplots showing genome-wide correlation between binding intensities (log2 CPM) of RAD21, SMC3, NIPBL and WAPL in WT (top) and *Brwd1^-/-^* (bottom) small pre-B cells. Datapoint color (purple) shows concurrent CTCF binding intensity. **h,** Total and coincident chromatin-bound peaks for static cohesin complexes (2F; RAD21+SMC3 without NIPBL+WAPL) in WT and *Brwd1^-/-^* small pre-B cells. **i,** Total and coincident binding peaks for dynamic cohesin complexes (4F; RAD21+SMC3+NIPBL+WAPL) in WT and *Brwd1^-/-^* small pre-B cells. **j**, Total and coincident binding of CTCF with dynamic cohesin (4F) in WT and *Brwd1^-/-^* small pre-B cells. **k**, Histograms demonstrating enrichment of RAD21, SMC3, NIPBL or WAPL at 5F (CTCF+RAD21+SMC3+NIPBL+WAPL)-bound sites in WT and *Brwd1^-/-^* small pre-B cells. **l,** Histogram of CTCF binding at 4F and 5F-bound chromatin sites described in (**h**) and (**j**) in WT and *Brwd1^-/-^* small pre-B cells. **m,** Histograms demonstrating enrichment of RAD21, SMC3, NIPBL and WAPL at 4F (RAD21+SMC3+NIPBL+WAPL)-bound sites in WT and *Brwd1^-/-^* small pre-B cells. **n,** Histogram demonstrating enrichment of BRWD1 at 4F and 5F-bound sites in WT small pre-B cells.

Peaks that contain only SMC3 and RAD21, and not NIPBL and WAPL which contribute to chromatin extrusion and cohesin recycling, were termed static or 2 factor binding (2F) cohesin. The number of static cohesin complexes were increased from 10,296 to 14,165 in *Brwd1^-/-^* small pre-B cells, and the distribution of static cohesin complexes in each cell population was very different (Fig. 3h). Chromatin-bound complexes containing SMC3, RAD21, NIPBL and WAPL (4 factor binding, 4F) were termed dynamic cohesion. These dynamic complexes were decreased from 20,475 in WT to 8,583 in *Brwd1^-/-^* small pre-B cells (Fig. 3i,j and Extended Data Fig.5b-i). This primarily reflected a loss of WT dynamic complexes in *Brwd1^-/-^* cells. These data suggest that BRWD1 regulates the equilibrium between chromatin-bound static and dynamic cohesin complexes.

Cohesin can bind at CTCF binding sites and either initiate loop extrusion or anchor TADs^19,32,33^. Examination of dynamic cohesin at CTCF-bound sites (5 factor binding, 5F) revealed variable complexes in *Brwd1^-/-^* cells with a slight increase in RAD21 binding and decreases in SMC3 and NIPBL compared to WT cells (Fig. 3k,l). Most WT dynamic cohesin complexes were not at CTCF sites, and these were preferentially lost in *Brwd1^-/-^* small pre-B cells (Fig. 3j). In contrast, there was a uniform and substantial decrease in all dynamic cohesin components that were not coincident with CTCF (4F) (Fig. 3m). BRWD1 was not coincident with 4F dynamic cohesin (Fig. 3n).

### BRWD1 converts static to dynamic cohesin complexes

In WT cells, most static cohesin complexes were not at CTCF sites (Fig. 4a). However, most of the static cohesin complexes gained in *Brwd1^-/-^* cells were at CTCF sites. Furthermore, there was enrichment of BRWD1 binding at both WT static cohesin/CTCF sites and at those CTCF sites that become bound by static cohesin in *Brwd1^-/-^* cells (Fig. 4b). Consistent with this, BRWD1 binding was enriched near CTCF anchors at TAD boundaries (Fig. 4c and Extended Data Fig. 6a). Overall, in the absence of BRWD1, there was an accumulation of static complexes at CTCF sites bound by BRWD1 in WT cells and a preferential loss of dynamic cohesin complexes from chromatin loops.

**Figure 4.**
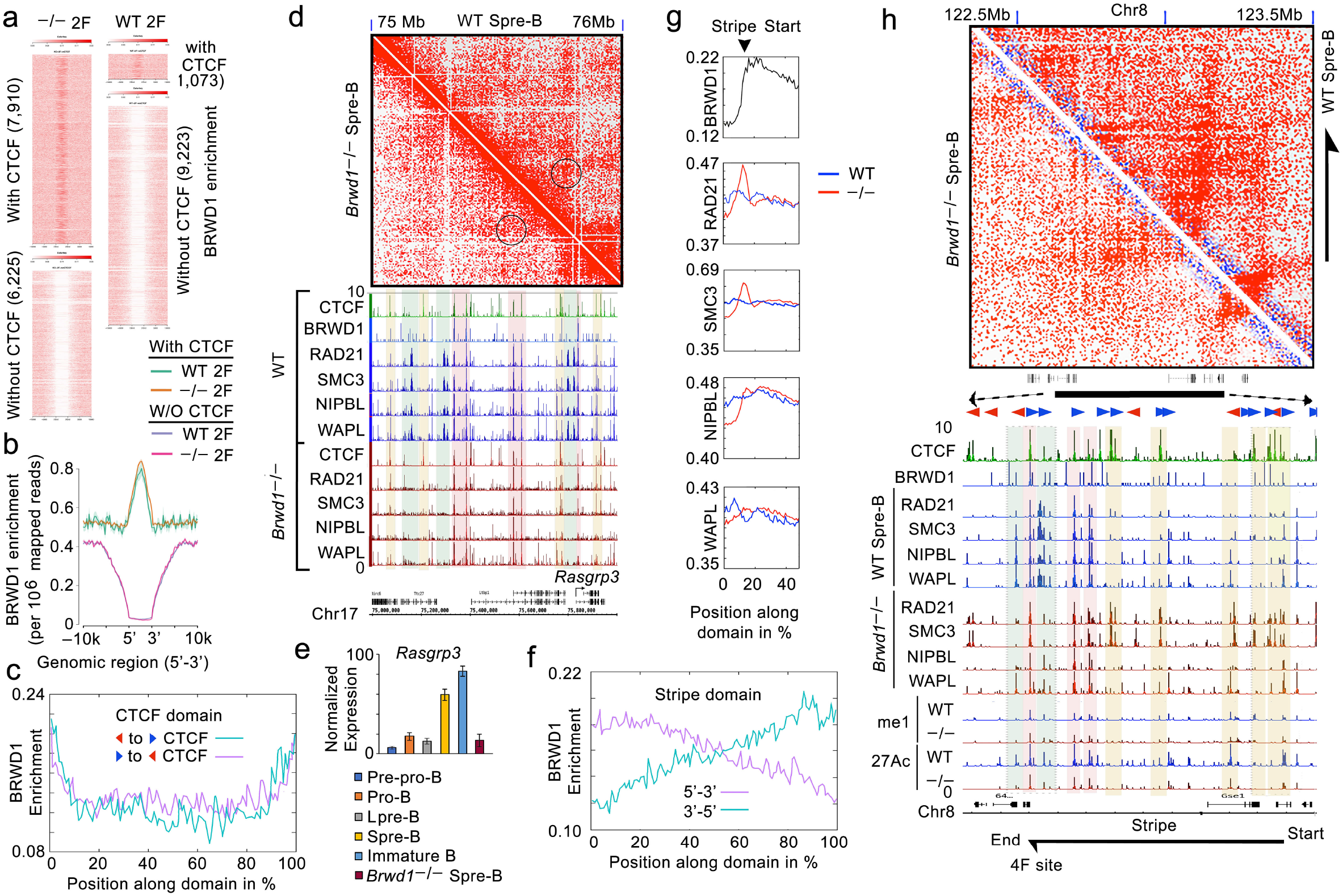
BRWD1 converts static to dynamic cohesin complexes. **a,** Heatmaps of BRWD1 tag density (WT) at static cohesin (2F) sites in *Brwd1^-/-^* and WT small pre-B cells organized by coincidence with CTCF. **b,** Histogram of BRWD1 enrichment at sites in **a**. **c,** Enrichment of BRWD1 across chromatin domains formed by convergent or divergent CTCF motifs. **d,** Hi-C interaction matrices of chromatin domain containing *Rasgrp3* in WT vs. *Brwd1^-/-^* small pre-B cells. *Rasgrp3* TAD contacts are circled. CTCF, RAD21, SMC3, NIPBL and WAPL chromatin binding are shown (bottom, all scaled 0-10 CPM). Yellow shaded boxes denote *de novo* static cohesion (2F) at CTCF sites in *Brwd1^-/-^* small pre-B cells. Green denotes *de novo* dynamic cohesion complexes (4F) in WT small pre-B cells. Pink denotes dynamic cohesion at CTCF sites (5F). **e,** Bar plot of *Rasgrp3* expression in indicated populations (RNA-seq, n=4). **F,** Enrichment of BRWD1 chromatin binding across 5’-3’ and 3’-5’ stripe domains in WT small pre-B cells. **G,** Normalized stripe domains demonstrating average distribution of bound BRWD1 (WT) and RAD21, SMC3, NIPBL and WAPL in WT and *Brwd1^-/-^* small pre-B cells. Strip origin is shown irrespective of orientation. Plots include flanking chromatin with stripe sites beginning at 16% and ending at 84% (not shown). **H,** Representative Hi-C contact matrices showing loss of a stripe in *Brwd1^-/-^* compared to WT small pre-B cells. Corresponding ChIP-seqs for CTCF and BRWD1 (WT), and then RAD21, SMC3, NIPBL, WAPL, H3K4me1 and H3K27Ac from WT and *Brwd1^-/-^* small pre-B cells. All ChIP-seq profiles plotted 0-10 CPM. Shaded areas are same as shown in “**d**”. Red and blue arrows indicate CTCF orientation.

An example of these binding distributions is afforded by the *Rasgrp3* locus (Fig. 4d-e). In WT cells, there was enhanced looping within the TAD, and this was associated with strong loading of dynamic cohesin complexes (green bars). In contrast, in *Brwd1^-^* cells, these dynamic complexes were absent and there were new static cohesin complexes proximal to both CTCF and WT BRWD1-bound sites (yellow bars). Cohesin complexes at CTCF sites were not substantially regulated by BRWD1 (pink bars). A multigeneic region of chromosome 5 (Extended Data Fig. 6b) and *Igk* (Extended Data Fig. 7a-c) provide other examples. Consistent with a lack of recombination in WT sorted small pre-B cells, cohesin complexes were readily detected across the *Igk* locus. These data suggest that BRWD1 converts static to dynamic cohesin complexes at TAD boundaries. Furthermore, our data demonstrate that cohesin complexes are heterogeneous in both function and regulation.

Stripes are features on Hi-C matrices indicative of asymmetrical cohesin loop extrusion and the juxtaposition of linear chromatin domains onto a single circumscribed region (Extended Data Fig. 7d). They often occur with promoter-enhancer interactions or during immunoglobin gene contraction^19,34–36^. BRWD1 binding at stripe loops was asymmetrical with enrichment at the origin, regardless of orientation (Fig. 4f). Likewise, RAD21 and SMC3 bindings were selectively enriched at the stripe origins in *Brwd1^-/-^* small pre-B cells (Fig. 4g). For example, at the chromosome 8 region containing *Irf8*, there was a loss of dynamic cohesin complexes (green bars) and an accumulation of static complexes at the stripe origin (yellow bars) co-incident with CTCF and BRWD1 in *Brwd1^-/-^* small pre-B cells (Fig. 4h). These data suggest that directed conversion of cohesin by BRWD1 at stripe origins dictate specific chromatin topologies.

### Cohesin conversion is common

Cohesin-mediated loop extrusion requires ATP^19,37^. We depleted WT small pre-B cells of ATP for two hours (Extended Data Fig. 8a-b) and repeated cohesin subunit ChIP-seq. ATP depletion redistributed cohesin component binding (Extended Data Fig. 8c-i). Compared to *Brwd1^-/-^* small pre-B cells, there was a further increase in static cohesin (2F) complexes (Fig. 5a-b) and loss of dynamic (4F) complexes (Fig. 5c-d). In ATP-depleted cells, static cohesin complexes strongly co-localized with CTCF-bound sites as did the relatively few persistent dynamic cohesin complexes. Furthermore, compared to WT and *Brwd1^/-^* small pre-B cells (Fig. 3d-g), there was a complete polarization of chromatin-bound cohesin complexes containing different components (Fig. 5e). Remarkably, there was a strong enrichment for BRWD1 binding at static cohesin sites regardless of CTCF co-incidence (Fig. 5f). In contrast, BRWD1 was not enriched at CTCF sites lacking static cohesin (Fig. 5g). At stripe origins, ATP depletion led to strong and specific enrichment for RAD21 and SMC3 (Fig. 5h). An example of this is the TAD-containing *Foxo1* locus (Fig. 5i). In both *Brwd1^-/-^* and WT ATP-depleted cells, there were loss of dynamic cohesin complexes and accumulations of static complexes at the stripe origin co-incident with CTCF and in proximity to BRWD1. These findings suggest that BRWD1 regulates the main mechanism of ATP-dependent cohesin remodeling in small pre-B cells.

**Figure 5.**
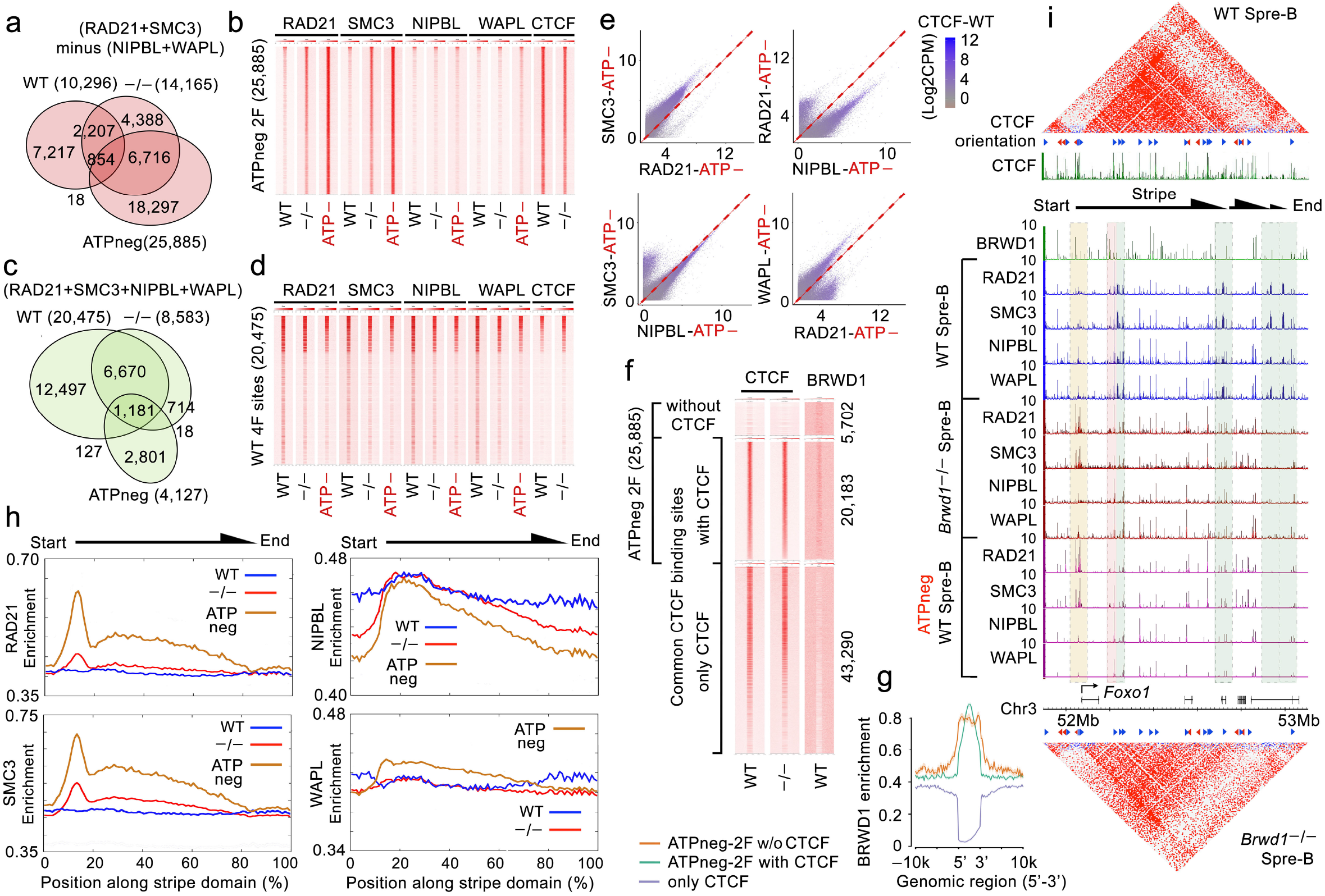
BRWD1-mediated cohesion conversion is a major ATP-dependent mechanism regulating cohesion function. **A,** Total and overlapping binding regions for static cohesion complexes (2F) in WT, *Brwd1^-/-^* and oligomycin A treated (ATP-depleted) WT small pre-B cells. **B,** Tag density heatmaps for RAD21, SMC3, NIPBL, WAPL and CTCF in WT, *Brwd1^-/-^* and ATP-depleted WT small pre-B cells at those 25,885 static cohesion sites (2F) observed in ATP-depleted cells (±10kb). **C,** Total and overlapping binding regions for dynamic cohesion complexes (4F) in WT, *Brwd1^-/-^* and ATP-depleted WT small pre-B cells. **D,** Tag density heatmaps for indicated ChIP-seq targets in WT, *Brwd1^-/-^* and ATP-depleted WT small pre-B cells at 20,475 4F sites detected in WT cells. **E**, Scatterplots demonstrating the genome-wide correlation between binding intensities of RAD21, SMC3, NIPBL and WAPL in ATP-depleted WT small pre-B cells (Log2CPM). Datapoint color (purple) is scaled to coincident CTCF binding intensity. **F,** Heatmaps of tag density for CTCF and BRWD1 (WT) at 25,885 static cohesion (2F) sites observed in ATP-depleted WT small pre-B cells with or without CTCF binding (top). This was compared to all other CTCF sites that were not coincident with static cohesion (bottom). **G**, Histogram showing BRWD1 enrichment at indicated regions as described in “**f”**. **h,**Normalized stripe domains demonstrating the average distribution of chromatin-bound RAD21, SMC3, NIPBL and WAPL in WT, *Brwd1^-/-^* and ATP-depleted WT small pre-B cells. Stripes are shown origin to end irrespective of orientation. Graphs include flanking chromatin with stripe sites beginning at 16% and ending at 84%. **I**, Hi-C interaction matrices of the *Foxo1* locus along with the corresponding binding profiles of CTCF (shown with motif orientation) and BRWD1 (WT) as well as RAD21, SMC3, NIPBL and WAPL in WT, *Brwd1^-/-^* and ATP-depleted WT small pre-B cells. Yellow shaded boxes denote *de novo* static cohesion (2F) at CTCF sites in *Brwd1^-/-^* small pre-B cells. Green denotes *de novo* dynamic cohesion complexes (4F) in WT small pre-B cells. Pink denotes dynamic cohesion at CTCF sites (5F).

## Discussion

These data indicate that BRWD1 induces and maintains 3D chromatin topology by converting static, chromatin-bound core-ring cohesin complexes at TAD boundaries and stripe origins to multimeric dynamic complexes that then dictate enhancer activation and mediate *Igk* locus contraction. Previous studies of cohesin regulation have focused on either modulation of cohesin subunit expression or mechanisms that target cohesin to chromatin including signaling pathways, transcription or transcription factors^23,24,38–40^. Our studies suggest that in small pre-B cells, cohesin conversion is the major ATP-dependent mechanism orchestrating changes in 3D chromatin topology required for cell state transition (Extended Data Fig. 9).

The striking enrichment of static cohesin at BRWD1-bound sites suggests that BRWD1 at CTCF sites directly converts static to dynamic cohesin. However, a more complicated conversion mechanism, in which BRWD1 independently regulates the removal of static complexes and the loading of dynamic cohesin, is possible. BRWD1 does not have predicted catalytic activity and therefore is unlikely to independently mediate cohesin conversion^41^. Rather, it is an approximately 263 kD scaffold protein that likely assembles effectors that direct ATP-dependent conversion of static to dynamic cohesin.

BRWD1 coordinately regulated chromatin looping, enhancer accessibility, and H3K27Ac. Furthermore, BRWD1 can regulate nucleosome positioning within open chromatin^4^. It remains to be determined if these functions are all a consequence of cohesin conversion or if BRWD1 integrates multiple pathways of gene activation. Furthermore, cohesin conversion does not explain BRWD1-mediated gene repression. These data suggest that other activities are encoded within the BRWD1 complex^42^. Further studies are needed to identify BRWD1-binding proteins and how different assemblies might enable different regulatory mechanisms of chromatin topology and gene transcription.

The expression and recruitment of BRWD1 to chromatin is coordinately regulated by the principal receptors of B lymphopoiesis, the IL-7 receptor, pre-BCR, and CXCR4^45,43^. This ensures that chromatin topological remodeling occurs in a precise developmental context. However, not all transcriptional changes in small pre-B cells can be ascribed to BRWD1^2^. Furthermore, in the immune system, the expression of BRWD1 is primarily restricted to B cells. Our data suggest that cohesin conversion is a major mechanism regulating gene topology. Therefore, it is likely that different lineage and stage restricted mechanisms regulate cohesin conversion to enable late B cell development and other specific cell state transitions.

## Supporting information

Extended Data Table 1. Raw reads and alignment of in situ Hi-C sequencing.

Extended Data Table 2. Raw reads and alignment of ChIP-seqs from WT and Brwd1-/- small pre-B cells.

Extended Data Table 3. Raw reads and alignment of ChIP-seqs from ATP-depleted WT small pre-B cells.

## Acknowledgements

We thank M. Olson and D. Leclerc for cell-sorting services and Pieter Faber for high throughput sequencing services. This work is supported by the US National Institutes of Health Grants R01 AI150860 (M.R.C. and M.M.) and R01 AI143778 (M.R.C.).

## Author Contributions

M.M. and M.R.C. designed the experiments; M.M. carried out and analyzed most of the experiments including the ChIP-seq; Y.H. prepared the Hi-C libraries. M.M-C. analyzed the high throughput sequencing and Hi-C data along with M.M. A.M. helped in initial analyses of Hi-C data and view-point analyses. M.L.V. helped in ATP depletion assay. M.L.V., N.E.W., M.K.O., Y.Y. and J.V. helped in genotyping, cell sorting and flow cytometry analyses. N.E.W. and K.G. helped in editing the manuscript. M.M. and M.R.C. oversaw the entire project and prepared the final draft of the manuscript.

## Data Availability

BRWD1 ChIP-seq and ATAC-seq data sets were deposited in the GEO database with accession number GSE63302. Hi-C data for pro-B cells can be obtained from GEO accession code GSE40173. Data for Hi-C of WT small pre-B, *Brwd1^-/-^* small pre-B and WT Immature B cells, ChIP-seq for H3K4me1, H3K27Ac, CTCF, RAD21, SMC3, NIPBL and WAPL from WT and *Brwd1^-/-^* small pre-B cells, and ChIP-seq for RAD21, SMC3, NIPBL and WAPL from ATP-depleted WT small pre-B cells were deposited in the GEO database with accession number GSE221519. The subseries for Hi-C is GSE221517. The subseries for ChIP-seq is GSE221518.

## Extended Data Figure Legends

**Extended Data Figure 1.**
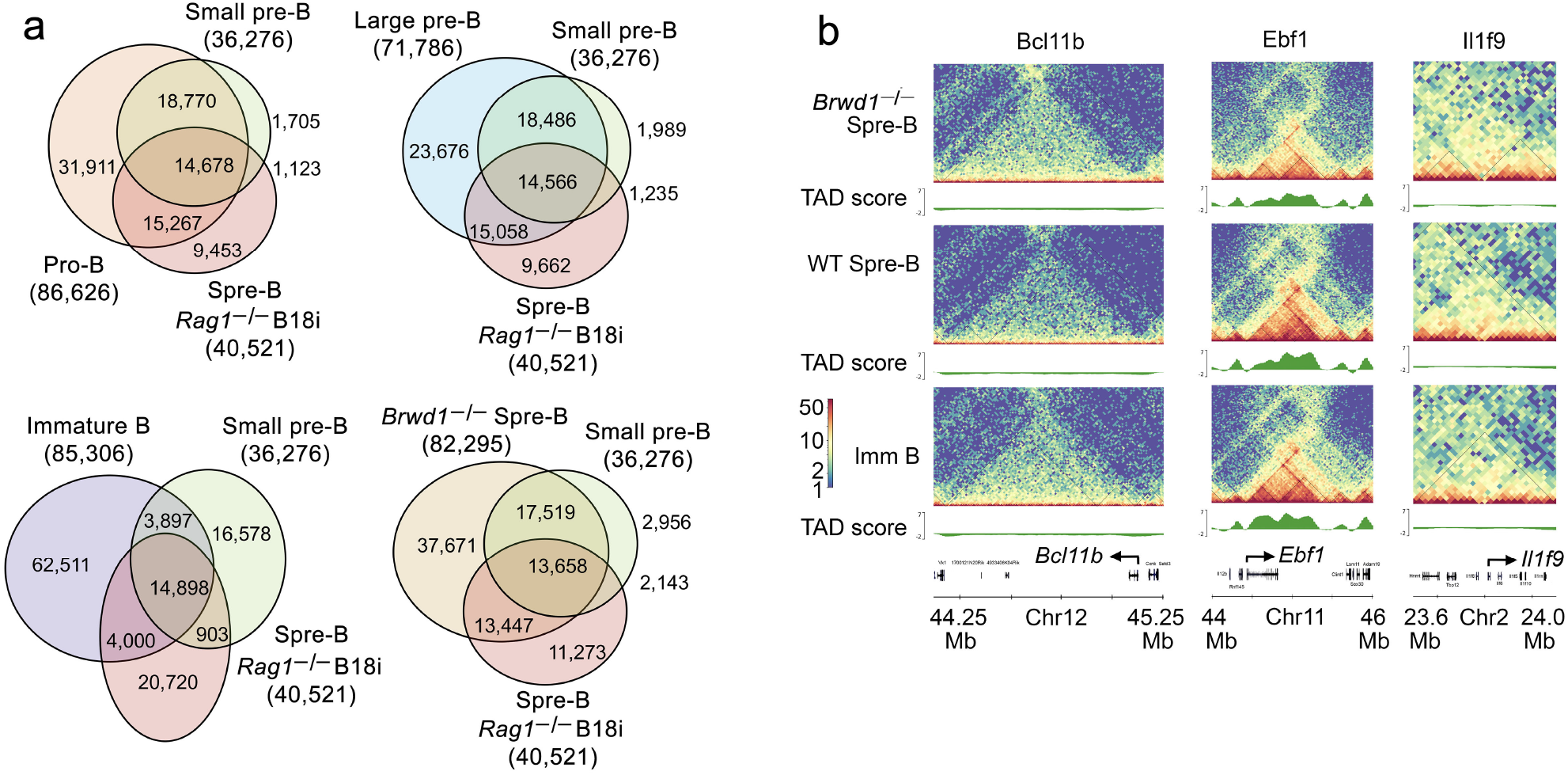
*S*mall pre-B cells have a unique developmental signature. **a,** Total and overlapping accessibility peaks (ATAC-seq) in WT pro-B, WT large pre-B, WT small pre-B, WT Immature B, *Brwd1^-/-^* small pre-B and *Rag1^-/-^*B18i (Igμ) small pre-B cells. **b,** Hi-C interaction matrices along with TAD scores in indicated cell populations for genomic regions containing T cell lineage specific (*Bcl11b*), B cell lineage specific (*Ebf1)* and granulocyte lineage specific (*Il1f9*) genes.

**Extended Data Figure 2.**
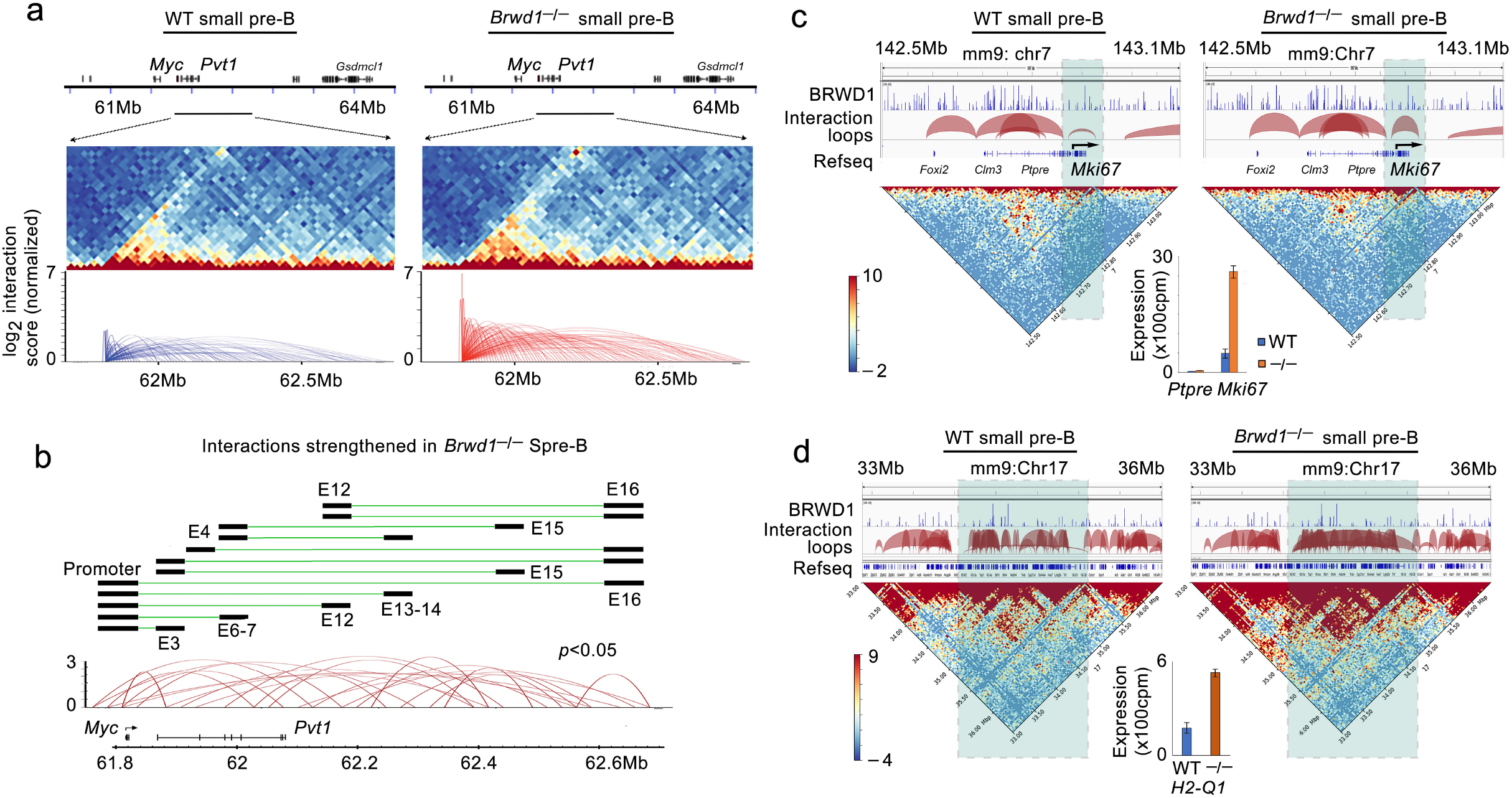
BRWD1 can repress chromatin interactions. **a**, Genomic region containing *Myc* in WT (left) and *Brwd1^-/-^* (right) small pre-B cells (top panels). Corresponding Hi-C interaction matrices of *Myc* locus (middle panels) and arc plots from point of view of *Myc* promoter region (bottom panels). The height of an arc represents the strength of interaction. **b,** Arc and line plots of significant interactions (*p*<0.05) strengthened in *Brwd1^-/-^* small pre-B cells compared to WT small pre-B cells. Schematic demonstrates promoter-enhancer and enhancer-enhancer interactions detected in *Brwd1^-/-^* small pre-B cells. **c,** *Mki67* gene region in WT small pre-B cells (left) and *Brwd1^-/-^* small pre-B cells (right). Corresponding Hi-C contact matrices and interaction loops as arc plots. The loop region containing *Mki67* is shaded. Expression of *Mki67* (RNA-seq; n=4) and *Ptpre* (gene located in adjacent loop) in WT and *Brwd1^-/-^* small pre-B cells shown as a bar plot (bottom). **d,** *H2-Q1* gene region shown and labeled as in “**c**”.

**Extended Data Figure 3.**
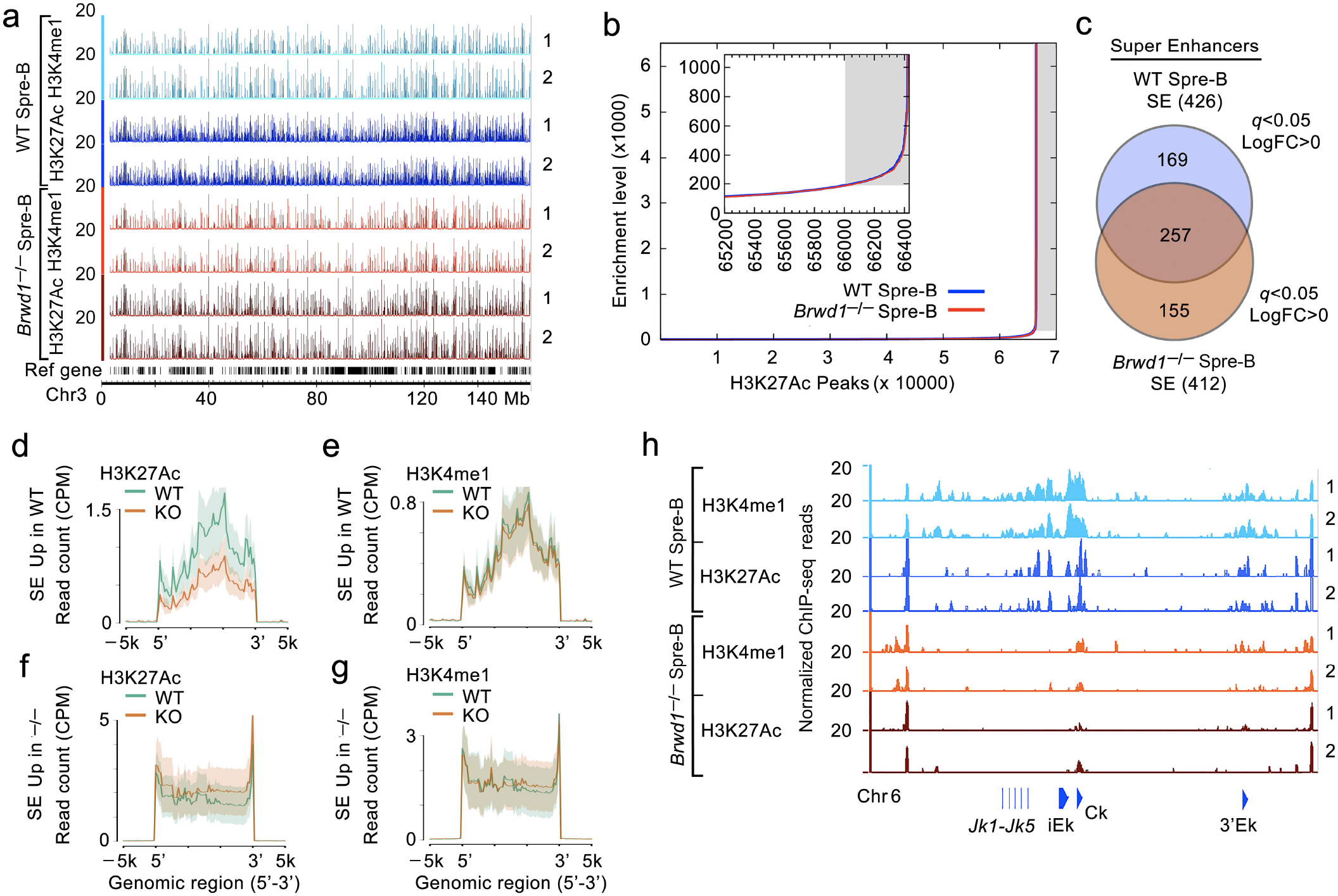
BRWD1 regulates super enhancers. **a,** ChIP-seq of H3K4me1 and H3K27Ac in WT and *Brwd1^-/-^* small pre-B cells across the entire chromosome 3. Data from replicates is shown. **b,** Identification of super enhancers (sEs) by enrichment of H3K27Ac binding. **c,** Venn diagram showing number of sEs in WT and *Brwd1^-/-^* small pre-B cells. **d-g**, Histograms showing enrichment of enhancer histone marks in sEs upregulated in WT or *Brwd1^-/-^* small pre-B cells. **h**, H3K4me1 and H3K27Ac binding in WT and *Brwd1^-/-^* small pre-B cells across the *Igk* locus from *Jk1-5*, through intronic enhancer (iEk) to 3’ kappa enhancer (3’Ek).

**Extended Data Figure 4.**
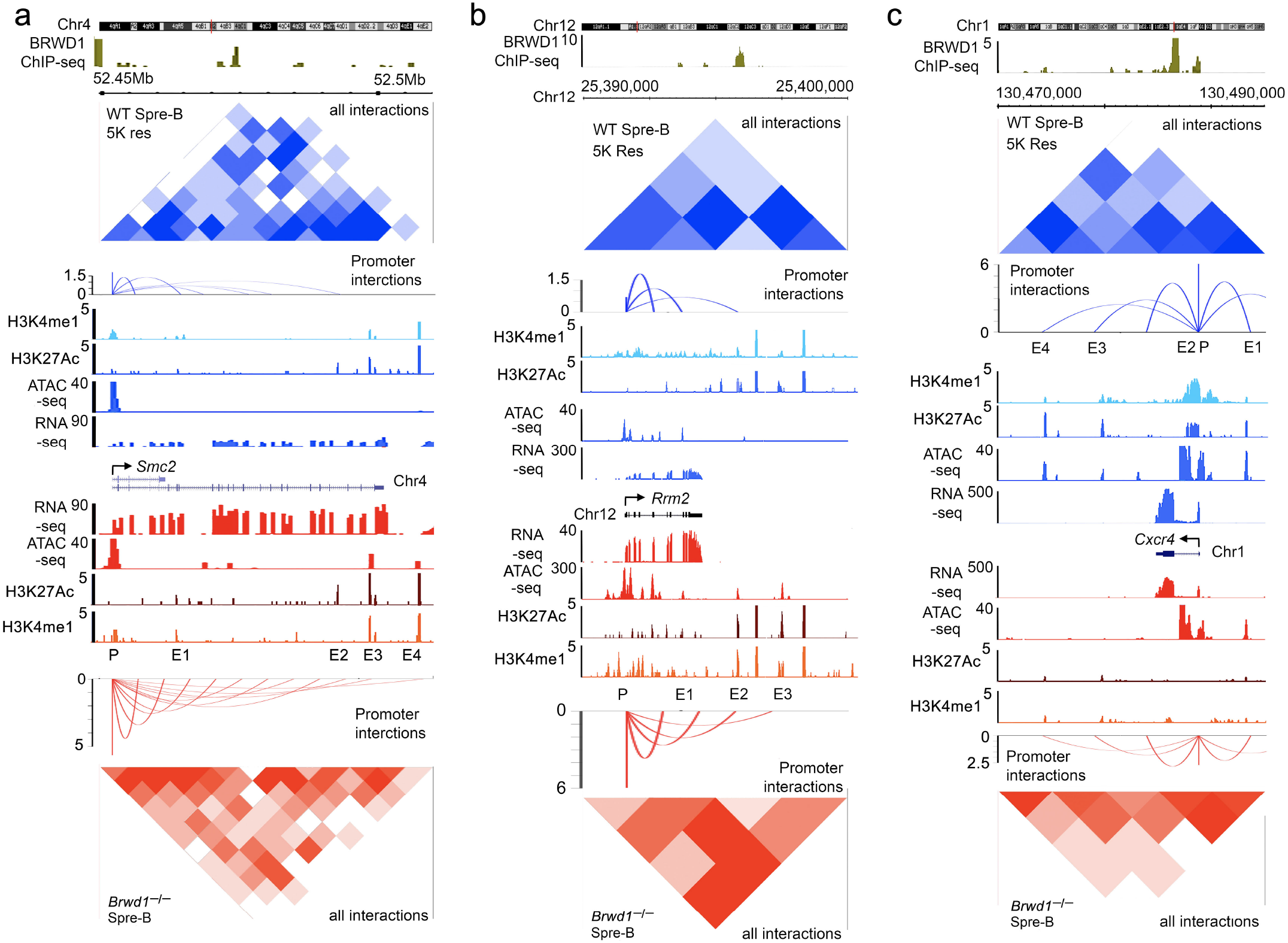
BRWD1 regulates enhancer-promoter interactions. **a-b,** BRWD1 repressed cell cycle related genes *Smc2* and *Rrm2*. **c,** BRWD1 induced gene *Cxcr4*. Corresponding Hi-C interaction matrices at 5 kb resolution, promoter-specific interactions as point-of-view arc plots, ChIP-seq for enhancer histone marks H3K4me1 and H3K27Ac, ATAC-seq and RNA-seq are shown. WT in upper blue and *Brwd1^-/-^* in lower red panels.

**Extended Data Figure 5.**
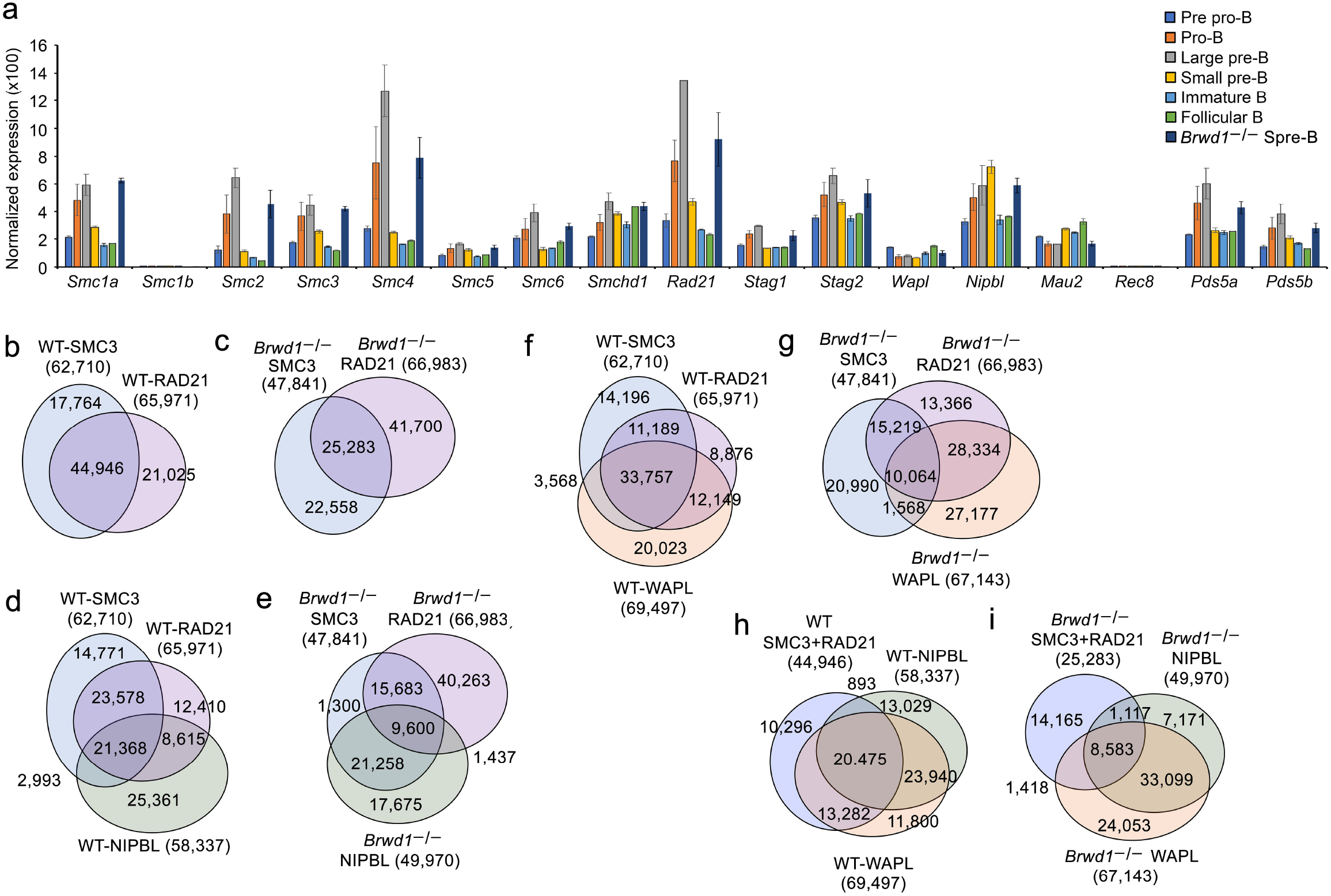
BRWD1 modulates cohesin binding. **a,**Bar diagram showing expression (RNA-seq; n=4) of indicated cohesin component genes in B cell progenitor subsets. **b-c,** Total and coincident genomic peaks for RAD21 and SMC3 in WT (**d**) and *Brwd1^-/-^*I) small pre-B cells. Total number of peaks for each population is shown in parentheses with number in each Venn region indicated. **d-e,**Total and overlapping coincident genomic peaks for RAD21, SMC3 and NIPBL in WT (**d**) and *Brwd1^-/-^* (**e**) small pre-B cells. **f-g,**Total and overlapping coincident genomic peaks for RAD21, SMC3 and WAPL in WT (**f**) and *Brwd1^-/-^* (**g**) small pre-B cells. **h-I,**Total and overlapping coincident genomic peaks for RAD21+SMC3 complex to NIPBL and WAPL in WT (**h**) and *Brwd1^-/-^* (**i**) small pre-B cells.

**Extended Data Figure 6.**
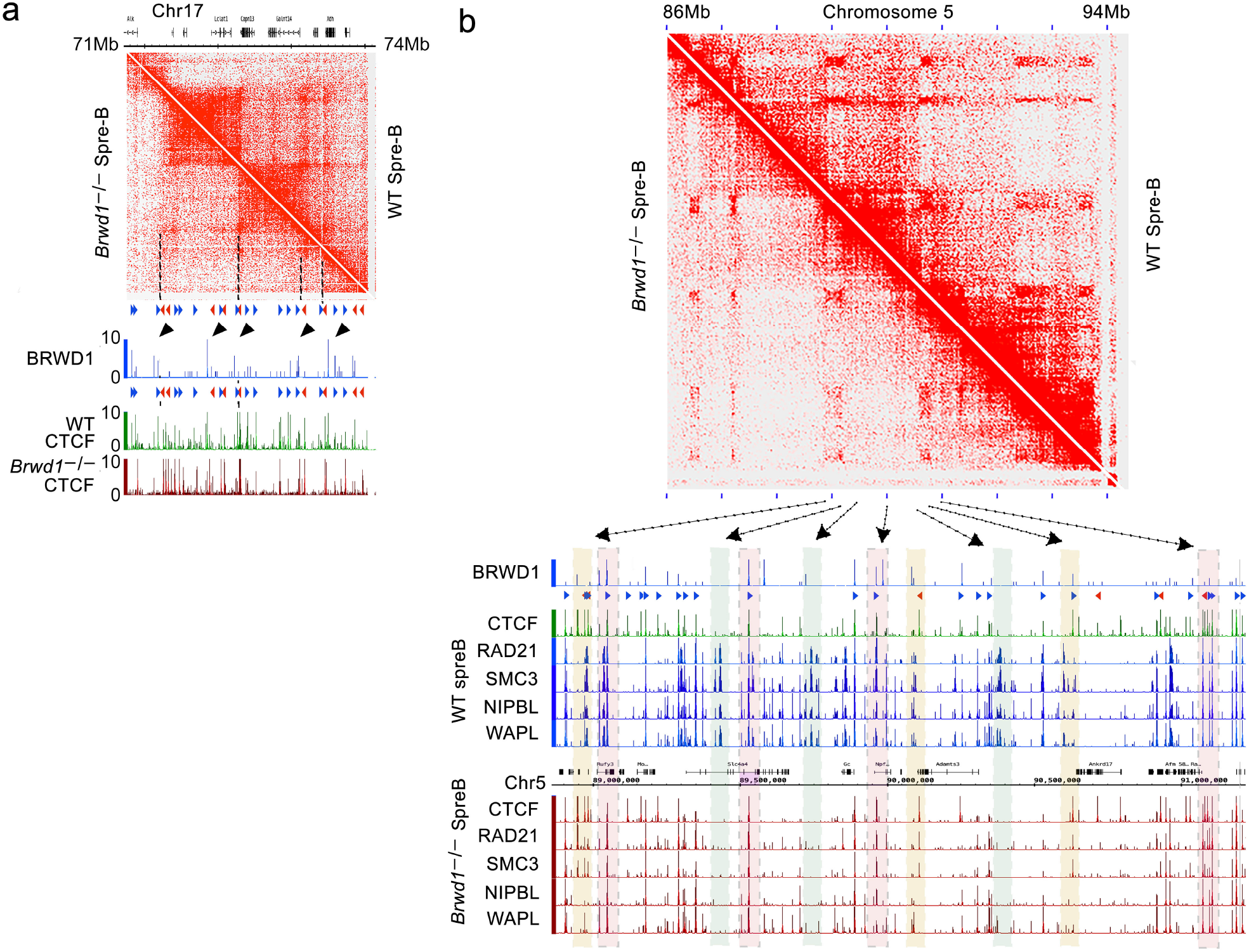
BRWD1 mediates static to dynamic cohesin conversion. **a,** Hi-C interaction matrix for indicated genomic region displaying BRWD1 binding near or at TAD anchor sites with CTCF binding (black arrowhead). Red and blue arrows indicate CTCF direction. **b,** Hi-C interaction matrix for indicated genomic region along with ChIP-seq profile of BRWD1 binding (WT), CTCF, RAD21, SMC3, NIPBL and WAPL binding in WT and *Brwd1^-/-^* small pre-B cells. Pink shaded boxes indicate regions with 5F (CTCF+RAD21+SMC3+NIPBL+WAPL; dynamic cohesin) binding in both WT and *Brwd1^-/-^* small pre-B cells. Yellow shaded boxes denote *de novo* static cohesin (2F) at CTCF sites in *Brwd1^-/-^* small pre-B cells. Green denotes *de novo* dynamic cohesin complexes (4F) in WT small pre-B cells. Pink denotes dynamic cohesin at CTCF sites (5F). All ChIP-seq profiles displayed as 0-10 CPM.

**Extended Data Figure 7.**
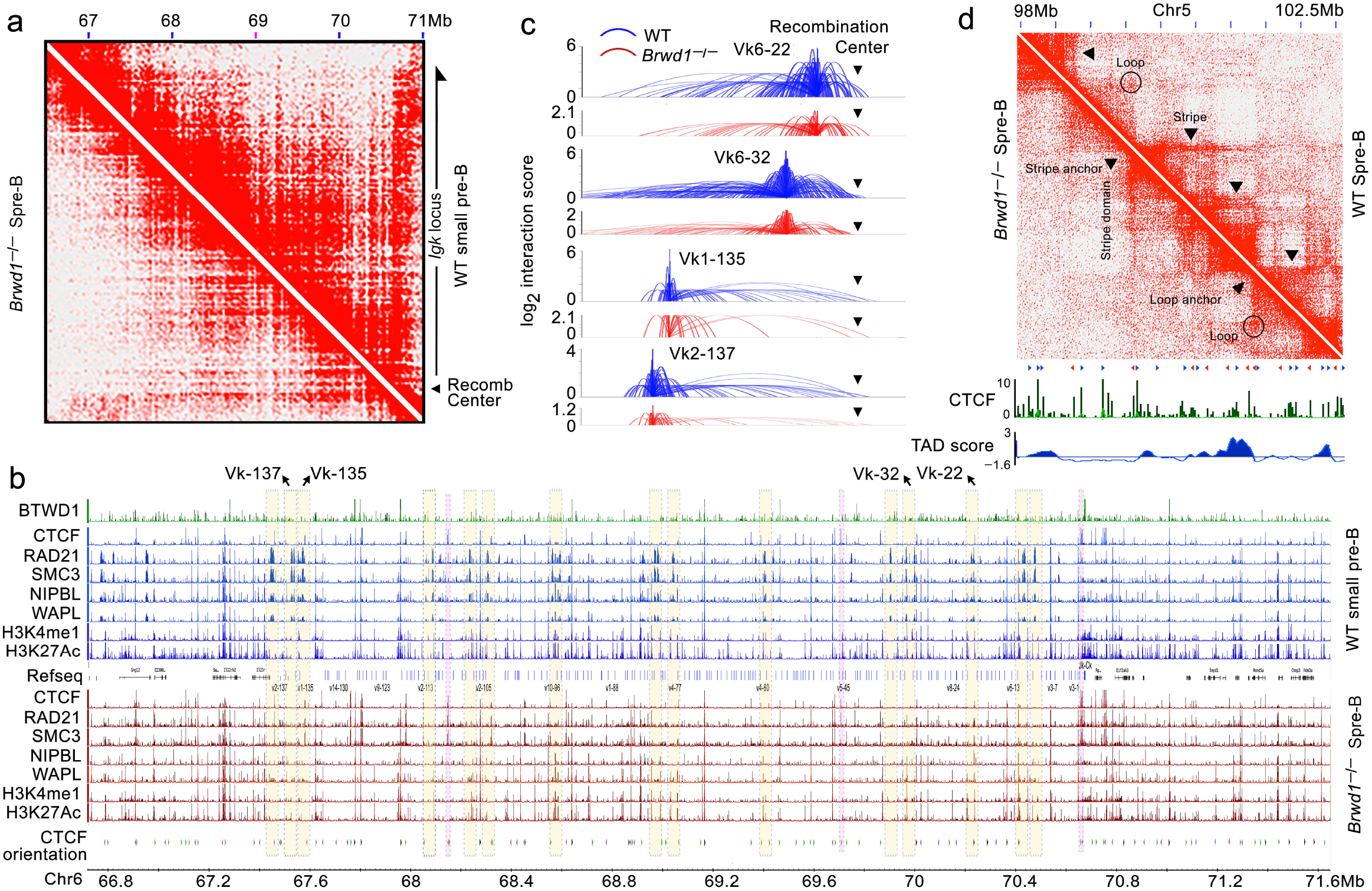
BRWD1 mediates loop extrusion at *Igk* locus by recruiting dynamic cohesin complexes. **a,** Hi-C interaction matrix for *Igk* locus demonstrating a stripe across entire WT *Igk* locus (~3.2 Mb, half arrow) that is anchored at the recombination center. **b,** ChIP-seq binding profile of BRWD1 (in WT), CTCF, RAD21, SMC3, NIPBL and WAPL along with histone marks H3K4me1 and H3K27Ac in WT and *Brwd1^-/-^* small pre-B cells across *Igk* locus. Yellow shaded boxes denote *de novo* static cohesin (2F) at CTCF sites in *Brwd1^-/-^* small pre-B cells. Green denotes *de novo* dynamic cohesin complexes (4F) in WT small pre-B cells. **c,** Point of view arc plots displaying normalized genomic interactions at indicated *Vk* gene segments in WT (blue) and *Brwd1^-/-^* small pre-B cells. Location of the recombination center is shown. **d,** Example showing stripes (black arrowheads) and conventional loop domains (circles) in WT and *Brwd1^-/-^* small pre-B cells (10 kb resolution) for a region of chromosome 5. The corresponding WT CTCF binding and TAD scores are displayed. Red and blue arrows indicate CTCF direction.

**Extended Data Figure 8.**
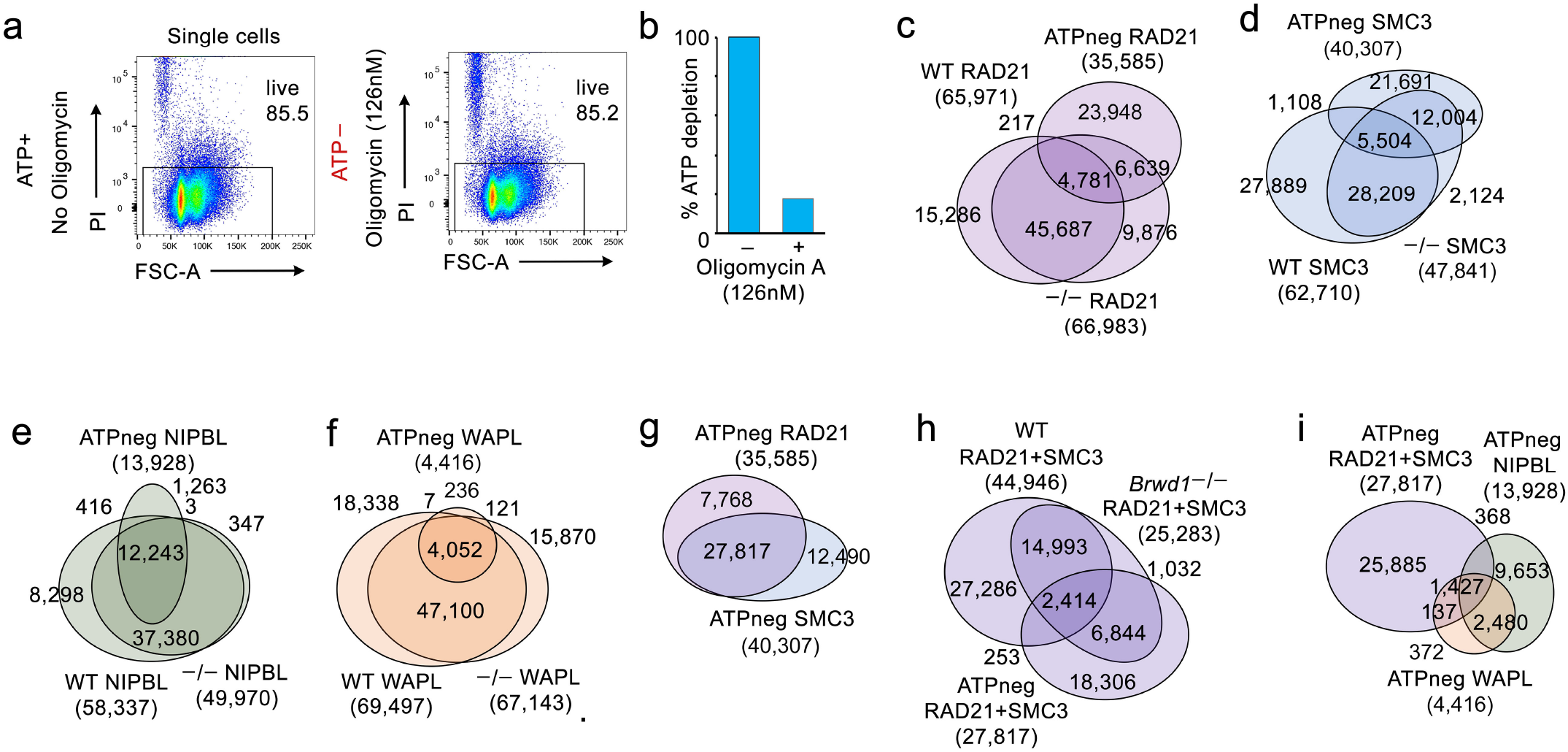
Cohesin recruitment in ATP-depleted WT small pre-B cells recapitulates BRWD1 deficiency. **a**, Flow cytometry demonstrating the percentage of live cells in WT small pre-B cells untreated (left) and treated with oligomycin A (126 nM) for 2 hours (right). **b**, Bar plot displaying ATP depletion (%) in WT small pre-B cells after 2 hours of treatment with oligomycin A (126 nM). **c-f,**Total and coincident binding peaks for RAD21 (**c**), SMC3 (**d**), NIPBL (**d**) and WAPL (**f**) in WT, *Brwd1^-/-^* and ATP-depleted WT small pre-B Cells. **g,** Total and coincident binding peaks for RAD21 and SMC3 in ATP-depleted WT small pre-B Cells. **h,** Total and coincident binding peaks for RAD21+SMC3 in WT, *Brwd1^-/-^* and ATP-depleted WT small pre-B Iells. **i,** Total and coincident binding peaks for RAD21+SMC3 with NIIPBL and WAPL in ATP-depleted small pre-B cells.

**Extended Data Figure 9.**
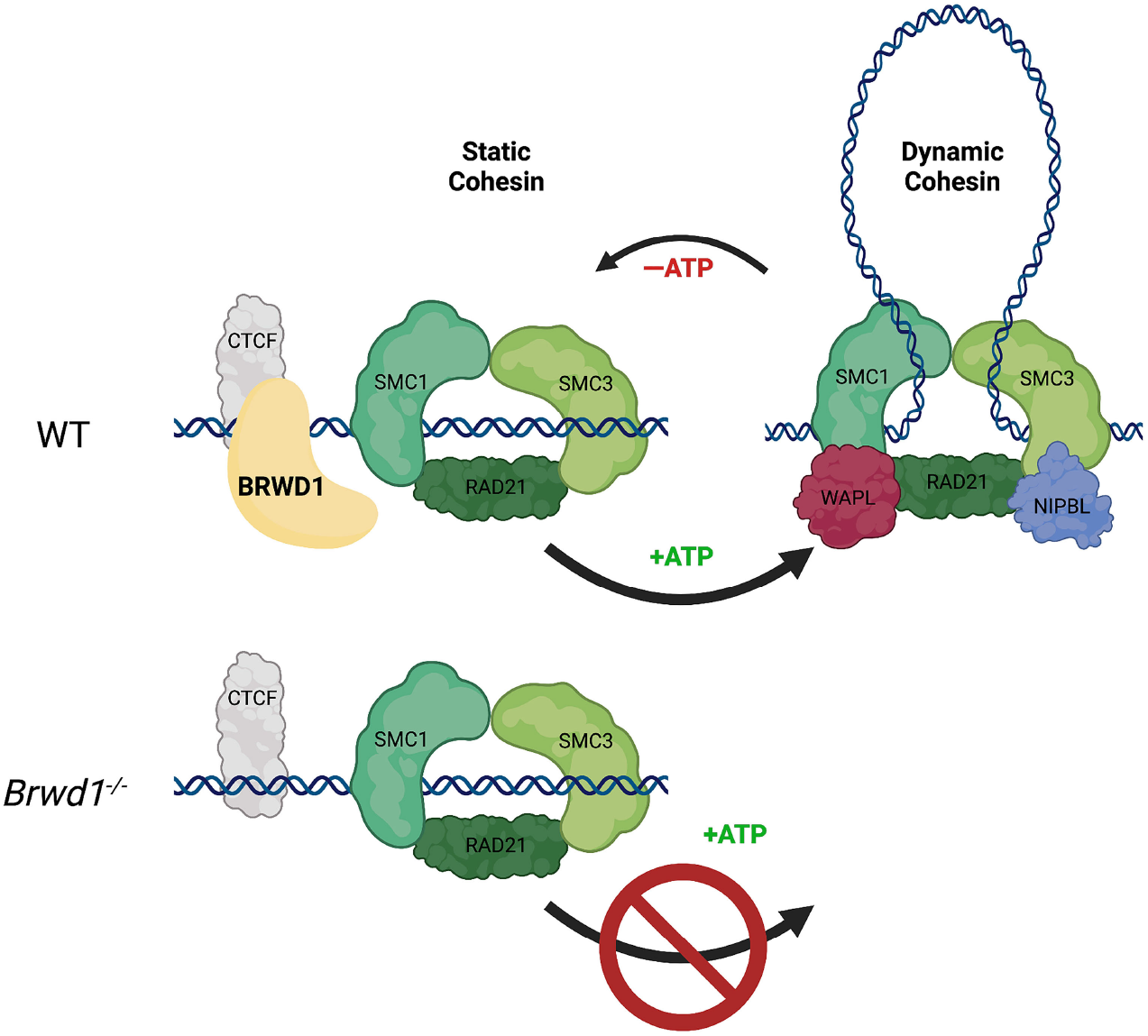
Model describing BRWD1-mediated static to dynamic cohesin conversion. BRWD1 complexes at CTCF sites convert chromatin-bound static cohesin to dynamic complexes competent for loop extrusion. At TAD boundaries this leads to a general increase in intra-TAD looping. In contrast, selective conversion at stripe origins induces directional loop extrusion and juxtaposition of linear chromatin regions onto origin proximate chromatin. In small pre-B cells cohesin conversion remodeled chromatin topology to mediate *Igk* contraction and dictate enhancer activation.

## Extended Data Tables

Extended Data Table 1. Raw reads and alignment of in situ Hi-C sequencing.

Extended Data Table 2. Raw reads and alignment of ChIP-seqs from WT and *Brwd1^-/-^* small pre-B cells.

Extended Data Table 3. Raw reads and alignment of ChIP-seqs from ATP-depleted WT small pre-B cells.

## Methods

### Mice

Wild-type (C57BL/6) and *Brwd1^-/-^* mice were housed in the animal facilities of the University of Chicago^2,4^. Mice were used at 6–12 weeks of age, and experiments were carried out in accordance with guidelines of the Institutional Animal Care and Use Committee of the University of Chicago.

### Isolation and flow cytometry of BM B cell progenitors

Bone marrow (BM) was collected from WT and *Brwd1^-/-^* mice as described previously^4^, and cells were resuspended in staining buffer (3% vol/vol FBS in PBS). Erythrocytes were lysed, and cells were stained with antibodies specific for CD19 (1D3, 1:400), B220 (RA3-6B2, 1:400), IgM (R6-60.2, 1:400) and CD43 (S7, 1:400) (all from BD Biosciences). Antibodies were directly coupled to fluorescein isothiocyanate, phycoerythrin, phycoerythrin-indotricarbocyamine, allophycocyanin, eFluor 450 or biotin. Pro-B cells (CD19+B220+CD43+IgM−), large pre-B cells (CD19+B220+CD43-IgM-FSChi), small pre-B cells (CD19+B220+CD43-IgM-FSClo), and immature B cells (CD19+B220+CD43-IgM+) were isolated by cell sorting with a FACSAria II (BD).

### Acute ATP depletion

For acute ATP depletion, small pre-B cells from WT BM were isolated as described above. Cells were placed at 37° C for 2 hrs in FACS buffer (1X PBS, 7.5% FBS) in the presence of Oligomycin A (126.4 nM, Sigma, #75351). ATP depletion was measured by the ATP Determination Kit (Thermo Fisher Scientific, #A2206) as described^19^, and cell viability was determined by flow cytometry.

### *In Situ* Hybridization and Immuno-FISH

*In situ* immuno-FISH was performed as previously described^8^.

### *In situ* Hi-C

*In situ* Hi-C was performed as previously described^6^. Briefly, WT and *Brwd1^-/-^* small pre-B and WT immature B cells (5×10^6^ for each stage) were crosslinked with 1% formaldehyde following a wash with ice cold PBS. Membranes were lysed to keep nuclei intact, followed by restriction digestion with 100U of MboI (NEB R0147). The DNA ends were marked with biotin, ligated proximally, and crosslinks were reversed. DNA shearing and size selection were then performed for fragments 300-500 bp. The amount of DNA was quantified with the Qubit dsDNA High Sensitivity Assay (Life technologies, Q32854), and biotin pull down was done to prepare the final *in-situ* Hi-C libraries (in duplicate). The number of reads and the alignments generated per sample are presented in Extended Data Table 1.

### Analysis of Hi-C sequencing

Hi-C processing was performed using Juicer ^44^ with default values, aligning to reference genome mm9; MboI fragment sizes were downloaded for mm9 from https://aryeelab.github.io/hichipper/RestrictionFragmentFiles/. Analysis was performed using the juicer.sh pipeline script. Contact matrices averaged over replicates were also computed by concatenating the duplicate-removed alignment files from each individual replicate and rerunning juicer_tools pre on the combined reads.

### Aggregate plots

To visualize promoter-enhancer interactions in WT and *Brwd1^-/-^* small pre-B cells, Hi-C aggregate plots were generated using hicAggregateContacts from HiCExplorer^47^ using 5. Kb resolution contact matrices with KR normalization. Analysis was performed between pairs of enhancer sites and the transcription start sites of active genes, and the mean observed/expected signal was plotted. The presence of an intense focal increase in the center indicates induced promoter contacts to the described enhancer groups.

### ChIP-sequencing

Chromatin from flow-sorted small pre-B cells (2×10^6^) from WT and *Brwd1^-/-^* mice were used for each ChIP experiment with anti-BRWD1 (E-15, Santa Cruz Biotechnology, sc-83517, lot J1508), anti-H3K4me1 (07-436, Millipore-Sigma), anti-H3K27Ac (07-360, Millipore-Sigma), anti-CTCF (Millipore Sigma, 07-729), anti-RAD21 (Abcam, ab154769), anti-SMC3 (Abcam, ab9263), anti-NIPBL (Proteintech, 18792-1-AP) and anti-WAPL (Proteintech, 16370-1-AP) antibodies. DNA libraries were prepared from sheared chromatin (200–600 bp). Libraries were sequenced on the Illumina Hiseq2500. The sequences were aligned to the mm9 reference genome (National Center for Biotechnology Information build mm9_NCBI_build_37.1) with BWA MEM^48^, and PCR duplicates were removed using Picard MarkDuplicates (Version 1.107).

### ChIP-seq peak calling and quantification

Peaks for ChIP-seq samples were called using MACS2 at a p-value threshold of 10^-5^ as done previously^4^. Normalized bedgraph tracks were generated using the SPMR flag and converted to bigwig using the UCSC tool bedGraphToBigWig. Peaks with a score >5 were retained. Normalized enrichment tracks for ChIP and input were stored as bigwigs.

Quantification and differential statistics steps were carried out separately for each ChIP-seq mark. First, peak calls from replicates in both WT and *Brwd1^-/-^* groups were merged using bed tools merge^49^. Peak and input enrichment counts in merged peaks was computed using featureCounts^50^. Input counts were subtracted from peak counts after adjusting for differences in sequencing depth for each library. Differential statistics (log2 fold-change and p-value) were computed using exactTest in edgeR^51^, and p-values were adjusted using the FDR correction.

### Differentially enriched peaks for histone marks and cohesin components

The set of reproducible peaks for histone marks and cohesin components was determined based on the peak enrichment statistics. First, normalized enrichment levels between replicates were compared, and peaks were retained based on (1) average CPM >5, and (2) squared difference in CPM over squared sum of CPMs <0.5 (i.e., high average enrichment and consistent enrichment between replicates). Adjacent peaks from this filtering process were then merged into a single region, especially to re-combine the “tiled” regions from the histone mark analysis. Enrichment scores for merged peaks were reported as the sum of the CPM values per peak divided by the length of the peak in kb. Then, both the replicates for individual WT and *Brwd1^-/-^* samples were merged and used for enrichment statistics. WT/*Brwd1^-/-^*-up peaks are differentially enriched (DE) peaks, based on the DE stats computed on merged peaks. DE peaks up or down were defined as Q < 0.05 and log_2_FC >1 (up by 2 fold) or < −1 (down by 2 fold). “Same” peaks were defined as Q > 0.2 and log_2_FC < 0.5 and > −0.5. They were segregated with the goal of having DE peaks be strongly up or strongly down and same peaks be clearly unchanged between groups. However, it is possible for some peaks to not appear in any of the DE sets if their DE statistics are in between these thresholds. These are peaks that show weak changes between groups.

### Super-enhancers

High-confidence peaks for H3K27Ac for each genotype were determined using Irreproducible Discovery Rate (IDR) analysis with an IDR threshold of 0.2. Peak calls within 12.5 kb were merged into putative super enhancers using bedtools merge^49^ and quantification of the enrichment of each ChIP and input sample was computed using featureCounts^50^. Input counts were subtracted from peak counts after adjusting for differences in sequencing depth for each library, and enrichment levels were averaged between replicates.

### Detection of stripes

Stripes were detected using a heuristic approach to match stripe enrichment patterns in Hi-C contact matrices, based on recent analytical approaches^19,34^. Observed/expected matrices with KR normalization were exported at 10 kb resolution. Row stripes, defined as stripes extending 3’ from the diagonal of the contact matrix, were computed first, and then matrices were transposed to search for column stripes, defined as stripes extending 5’ from the diagonal, using the same procedure.

For row stripes, first a signal over background was computed for each row over all genomic bins within 3 Mb of the diagonal. The signal for each bin along the row was computed as the median value of all bins within 2 indices above and below and to the left and right (e.g., at 10kb resolution, a 50kb x 50kb square centered on the bin). Background for each bin was computed as the median value over all bins within 6 but greater than 2 indices (e.g., at 10kb resolution a 130kb x 130kb square, with the middle 50kb x 50kb square excluded). Signal over background for the bin was computed as the difference between signal and background values. Values were then converted to a binary sequence: values > 0.2 were set to 1, other values set to 0.

Candidate stripe patterns in the binary sequence were determined by extending along the row from the diagonal as long as the following criteria were met: (1) average number of positive bins in the stripe ≥ 0.6, (2) average number of positive bins in the right-most 20 bins ≥ 0.6. Stripes longer than 9 bins were reported.

### Differential analysis of stripes

Stripes were determined first in the replicate-averaged contact matrices for better power. To compare stripe values between replicates and enable quantitative comparisons between genotypes, we first combined stripes between WT and *Brwd1^-/-^*, merging overlapping or adjacent stripes. Overlapping stripes were those within the same row. Adjacent stripes were defined as those within 10 kb above or below for row stripes, or within 10 kb left or right for column stripes. Merged stripe coordinates were computed as the average of the coordinates for all stripes that were combined. Each combined stripe was quantified from the contact matrix (10 kb resolution, observed/expected, KR normalization) for each individual replicate as the trimmed mean of all bins within the stripe. Differential stripes were identified using limma^45^, adjusting p-values using the FDR correction.

### Analysis of the directionality of CTCF at stripe anchors

We obtained CTCF motif directionality for CTCF peaks identified in the WT and *Brwd1^-/-^* small pre-B cells as described previously^19^. Directionality of CTCF peaks at stripe anchors was considered only when all peaks overlapping the anchor were attributed to a CTCF motif.

### Stripe footprinting

Stripe domains were expanded by 20% on either end and stored as a bed file; row stripes were considered plus strand and column stripes minus strand. Enrichment values for ChIP marks over each stripe were extracted from bigWig files using bigWigToBedGraph^52^, reversed for minus strand (column) stripes, and the stripe size was rescaled to fit on a 0-100 scale. Average enrichment values were computed over 100 equally sized bins on this scale. Trimmed mean value of every bin was computed over all stripes to generate an average footprint that is representative of enrichment levels over the body of a stripe.

### View-point Plots

Hi-C matrices were jointly normalized using the hic_loess function from HiCompare^53^, and difference detection was performed using the hic_compare function. The plotBedpe function from Sushi^54^ was used to plot differences with a p-value less than 0.05 or 0.01 unless otherwise mentioned.

